# Simulating Multi-Colour Single-Molecule Localisation Microscopy Using an RGB Camera

**DOI:** 10.64898/2026.04.15.718692

**Authors:** Ava E D Kelly, John S H Danial

## Abstract

High-order multiplexing in single-molecule localisation microscopy (SMLM) is limited by trade-offs between spectral discrimination, imaging speed, and experimental complexity. Here, we show that RGB cameras provide a simple and scalable solution for multi-colour SMLM by exploiting their intrinsic spectral sensitivity for statistical fluorophore discrimination. Using a realistic simulation framework incorporating experimentally derived photon budgets, optical response functions, and camera noise, we achieve simultaneous classification of up to six fluorophores with a mean precision of ∼98%, including perfect discrimination of spectrally overlapping dye pairs, while maintaining an average localisation precision of ∼3.2 nm. Performance remains robust to variations in classification thresholds but degrades with increasing fluorophore number and reduced photon budgets due to spectral overlap and photon noise. These results establish RGB detection as a cost-effective and experimentally straightforward alternative to conventional spectral imaging approaches, enabling accessible, high-throughput multiplexed super-resolution imaging.

## Introduction

Single-Molecule Localisation Microscopy (SMLM) has revolutionised biological imaging by enabling the organisation of multi-protein complexes to be resolved at molecular resolutions, directly in situ^1,2^. This capability has made it possible to visualise flexible assemblies such as clathrin-coated pits^3^, filamentous proteins^4^, and apoptotic complexes^5^ that would typically evade structural characterisation using alternative techniques, allowing them to be imaged in their native environments. Despite substantial advances in improving SMLM resolution, achieving rapid, accessible, and high-order multi-target imaging remains a significant challenge.

On one hand, spectral imaging approaches such as ratiometric imaging using beam-splitting devices are relatively easy to integrate into a wide range of custom and commercial instruments. However, they are typically limited to multiplexing up to three targets and often require reduced fields of view (FoVs)^6^. Diffraction gratings^7^ and prisms^8,9^ can help overcome this limitation, but at the cost of significantly increased experimental complexity, restricting their use to specialist laboratories.

While it is in principle possible to combine ratiometric and multidimensional imaging using multiple dichroic beam splitters and cameras that scale with the number of targets, such configurations, though recently demonstrated^10^, are both economically and technically challenging to implement.

On the other hand, sequential imaging strategies based on orthogonal chemistries such as DNA Point Accumulation for Imaging in Nanoscale Topography (DNA-PAINT)^11^ and its variants, including secondary label-based unlimited multiplexed DNA-PAINT (SUM-PAINT)^12^ and fluorogenic labelling with transient adapter-mediated switching for high-throughput DNA-PAINT (FLASH-PAINT)^13^, as well as DNA or protein barcoding^14,15^ and multiplexing using erasable signals^16^, can enable imaging of up to 30 targets within a single FoV. However, these approaches require intricate DNA sequence design where applicable, complex and labour-intensive experimental protocols, and, importantly, suffer from markedly reduced imaging speeds.

More recently, intrinsic photophysical properties of fluorophores such as intensity^17^ and fluorescence lifetime^18^ have been exploited to distinguish between different labels. Intensity-based discrimination represents the simplest route to rapid, multi-colour SMLM, but has so far only been demonstrated using three red-emitting dyes. Lifetime-based approaches, while capable of distinguishing up to eight fluorophores, have been limited to confocal implementations and confocal-type super-resolution modalities rather than wide-field imaging.

Inspired by the human visual system, specifically the role of cone cells in enabling accurate colour discrimination, we simulate the performance of RGB cameras for high-throughput, full-colour SMLM. Unlike conventional monochrome sensors, RGB cameras retain sufficient spectral information to enable statistical discrimination between fluorophores, even when their emission spectra overlap, effectively introducing colour as an additional measurement dimension. Our simulations demonstrate up to 98% accurate classification of six fluorophores imaged simultaneously, paving the way for the experimental realisation of straightforward, high-order, multi-colour SMLM at full imaging throughput.

## Methods

Nine commercially available fluorophores were included in the simulations, with emission maxima ranging from 519 nm (AF488) to 682 nm (CF660R) and experimentally measured photon outputs per 100 ms ranging from 2073 to 23195 photons (**Supplementary Information**). Photon outputs accounted for differences in fluorophore absorption, given that the excitation source was fixed at 488, 560, or 642 nm, and were derived from experimental measurements of total fluorescence prior to photobleaching^19^. These fluorophores were chosen to span the spectral range accessible with the three excitation wavelengths and to provide a representative set of fluorophores commonly used in multicolour super-resolution experiments.

A virtual microscope was simulated in Python to replicate a typical experimental setup. The optical configuration consisted of a three-colour excitation source, which was directed through a 100×, 1.25 numerical aperture objective lens. Emitted fluorescence was collected by the same objective, passed through a penta-band dichroic mirror (DI01-R405/488/561/635/800, Semrock) and a quad-band emission filter (FF01-446/523/600/677, Semrock), and finally focused via a 70 mm tube lens onto an RGB camera (Blackfly S BFS-U3-32S4C-BD, FLIR) to yield a final pixel size of 98.6 nm. The spectral response of each component, including the excitation sources, dichroic, emission filter, and camera quantum efficiency, was measured experimentally and combined with each fluorophore’s emission spectrum to generate a single compounded spectral response curve for each fluorophore. This curve determined the fraction of photons detected in each of the red, green, and blue channels of the RGB camera for a given fluorophore.

Photon emission from each fluorophore was then sampled from a Poisson distribution to match experimentally observed photon statistics^20^, and the emission pattern was convolved with a Gaussian point spread function defined by the objective numerical aperture and the fluorophore emission wavelength. The resulting photon distributions were converted to electron counts and digitised into Analog-to-Digital Units (ADUs) using experimentally measured camera parameters, including baseline offset, dark current, read noise, thermal noise, and gain, with additional shot noise added to replicate realistic measurement conditions^20^. Signals were then distributed across the RGB channels according to the compounded spectral response for each fluorophore, creating simulated images that closely matched those obtained in physical experiments.

For each simulated emitter, mean red, green, and blue pixel intensities were extracted from a 5 × 5-pixel region centred on the fluorophore, and histograms of these intensities were compiled to characterise the expected intensity distributions in each colour channel. Identification regions were defined for each fluorophore to encompass 60% of the intensity values, although this fraction could be adjusted to optimise either specificity or sensitivity. An emitter was assigned to a fluorophore if its mean R and G intensities fell exclusively within the identification regions of that fluorophore; emitters falling outside all identification regions or within the identification regions for more than one fluorophore were labelled as unknown.

To assess localisation performance, the simulated images were further processed using the ImageJ plugin ThunderSTORM^21^. A wavelet filter was applied with B-Spline order 3 and B-Spline scale 2. Approximate localisation of molecules was performed via local maximum detection, using an intensity threshold equal to the standard deviation of the filtered image. Sub-pixel localisation was performed by maximum likelihood fitting of a Gaussian, with a fitting radius of 3 pixels and an initial sigma of 1 pixel. Resulting localisations were filtered to retain PSF widths between 90 and 140 nm, and further filtered by intensity and localisation precision, with thresholds adjusted per image to remove outliers and signals arising from overlapping localisations.

## Results

Using experimentally determined photon budgets reported previously^19^ (mean photon count of 8,543 for 100 ms exposures, **Supplementary Information**) and defining an identification region spanning 60% of the spectral space, we first evaluated the performance of our classification framework across six dyes: Alexa Fluor 488 (AF488), Alexa Fluor 647 (AF647), ATTO 488 (AT488), Cy3B, CF660R, and JF585. Under these conditions, the system achieved a high mean classification precision of approximately 98%. Notably, four dyes (AF488, AT488, Cy3B, and JF585) were classified with 100% precision, indicating near-perfect separability within the defined identification space (**Supplementary Information**).

In parallel, we quantified the localisation precision achievable using an RGB camera under the same experimental constraints. Across all dyes, an average localisation precision of 3.19 nm was obtained. Among these, Cy3B which is commonly employed in DNA-PAINT experiments exhibited particularly high performance, achieving a localisation precision of 1.78 nm, consistent with its favourable photophysical properties (**Supplementary Information**).

Having established baseline performance under realistic conditions, we next systematically challenged the framework by varying two key parameters: (i) the width of the identification region (ranging from 20% to 80%), and (ii) the number of distinguishable dyes (increasing from six to nine). Varying the identification region width had a relatively minor impact on classification precision, which decreased only modestly from 99.67% to 95.53%. However, this came at the expense of a substantial increase in the abstention rate (defined as the fraction of “unknown” classifications), which approached 95.34% as the identification region was narrowed to 20% (**Supplementary Information**). This behaviour reflects a more conservative classification regime, in which the system increasingly rejects ambiguous detections rather than risking misclassification.

In contrast, increasing the number of dyes produced the opposite effect. While the abstention rate remained relatively stable at approximately 60% (ranged from 60.50% to 66.85%), the classification precision decreased significantly to 85.19%, indicating increased overlap in the spectral signatures and a higher likelihood of incorrect assignments when more dyes are included (**Supplementary Information**).

Finally, we investigated the impact of reduced photon budgets to assess the feasibility of using RGB-based classification and localisation in photon-limited applications, such as single-molecule tracking, molecular counting, and Förster resonance energy transfer (FRET). A five-fold reduction in the mean photon budget resulted in a moderate increase in the abstention rate (from 60.50% to 68.64%), but a more pronounced degradation in classification precision, which dropped from 97.91% to 73.70% (**Supplementary Information**). In parallel, localisation precision deteriorated from 3.19 nm to 5.77 nm (**Supplementary Information**).

## Conclusion

Although RGBW cameras have previously been demonstrated for two-colour SMLM^22^, we show here for the first time, through computational analysis, that RGB cameras enable high classification precision for 6–9 dyes imaged simultaneously, achieving performance comparable to, and in some cases exceeding, that of more sophisticated spectral discrimination systems used in SMLM. Notably, even spectrally proximate dye pairs such as ATTO 488 and Alexa Fluor 488, or Cy3B and JF585, can be discriminated with 100% precision, highlighting the resolving capability of this approach despite the limited spectral information available from RGB detection.

Under low photon budget conditions, classification precision decreases, as expected. This reduction can be partially mitigated by limiting the number of dyes included in the analysis. At low photon counts, particularly for dim fluorophores, a substantial fraction of simulated emitters is rejected during pre-processing (**Supplementary Information**). This effect arises from an apparent broadening of the point spread function, which occurs when emitter intensities approach the camera noise floor and reduces both detectability and fitting reliability. These observations highlight the sensitivity of both classification and localisation performance to photon statistics, while also demonstrating that meaningful performance can still be achieved under photon-limited conditions.

Although the simulations capture many aspects of a realistic experimental system, certain optical aberrations, including spherical and chromatic aberrations, were not explicitly modelled. This simplification is justified by the high correction quality of modern objective and tube lens systems, which minimise such aberrations under typical imaging conditions. In addition, while the simulations were performed using a commonly employed combination of dichroic and emission filters for multi-colour single-molecule experiments, alternative filter sets may further improve both classification and localisation precision.

Finally, we demonstrate that high-fidelity multi-colour SMLM can be achieved using an industrial RGB CMOS camera, providing a cost-effective alternative to conventional detection systems. Further improvements in localisation precision are expected with scientific-grade RGB CMOS cameras, which offer higher quantum efficiency and lower noise characteristics.

**Figure 1.**
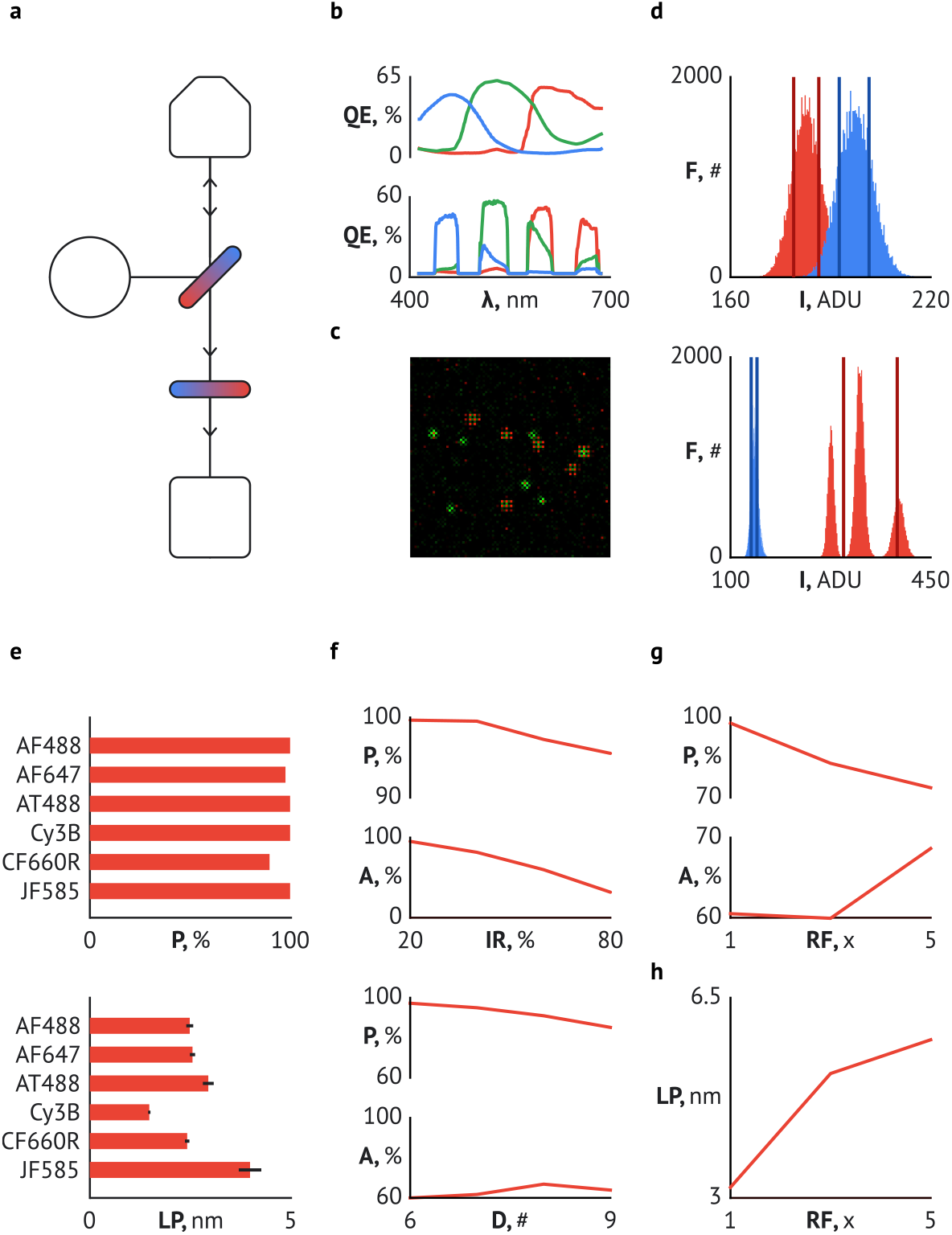
**a**, A 488 nm, 561 nm, and 638 nm laser excites a sample. Emission is separated from excitation using a dichroic filter (DI01-R405/488/561/635/850, Semrock) and an emission filter (FF01-446/523/600/677, Semrock) before imaging on an RGB camera (BlackflyFS-U3-32S4C-BDIR). The objective (100x) is paired up with a 70 mm tube lens to produce a final pixel size of 98.6 nm. **b**, The averaged spectral response of the blue, green, and red pixels of the camera (top) is compounded with the spectral response of the dichroic and emission filters (bottom). **c**, Single emitters are convoluted with a Gaussian Point Spread Function (PSF) whose width is calculated using each fluorophore’s emission wavelength and the objective’s numerical aperture. Example image showing single Atto488 and Alexa 647 dyes simulated on a camera with an RGB Bayer filter. **d**, The RGB intensity of each emitter is measured and plotted as histograms. An identification region (IR) is defined to include 60% of the intensities. An unknown emitter is assigned to a fluorophore if its RGB values fall within that fluorophore’s IR; otherwise, it is rejected. Intensities (I) of Atto488 (blue) and Alexa 647 (red) and their respective IRs are plotted in the green (top) and red (bottom) channels. **e**, Classification precision (P), calculated as the number of correct identifications (C) divided by the sum of C and the number of incorrect identifications (I), is quantified for 6 dyes (top) with realistic photon budgets extracted from^19^ for DNA-PAINT experiments. Our results indicate an average classification precision of 97.9%. Furthermore, the localisation precision (LP, bottom) is not worsened using an RGB camera, where Cy3B is localisable with a precision of 1.78 nm. **f**, P and the abstention rate (A), calculated as unknown identifications (U) divided by C + I + U, quantified against width of IR (top) and number of dyes (D below). **g**, P and A quantified against the average photon budget reduction factor (RF), where 1 represents an average of 8,543 photons and 5 represents an average of 1,709 photons. **h**, LP quantified against RF.

## Data availability

All simulated datasets are available on request.

## Code availability

All codes are publicly available on https://github.com/aedk02/smlm-RGB-simulator.

## Author contributions

JSHD conceived the study. AK performed all simulations with assistance from JSHD. JSHD and AK wrote the manuscript.

## Acknowledgments

JSHD acknowledges pump-priming funding from the RS MacDonald Charitable Trust, a collaborative research grant from the Royal Society of Edinburgh (RSE), and a Medical Research Council (MRC) Career Development Award (UKRI1421). AK acknowledges an EPSRC PhD studentship.

