## Supplementary Information for "Simulating Multi-Colour Single-Molecule Localisation Microscopy Using an RGB Camera"

### Figures

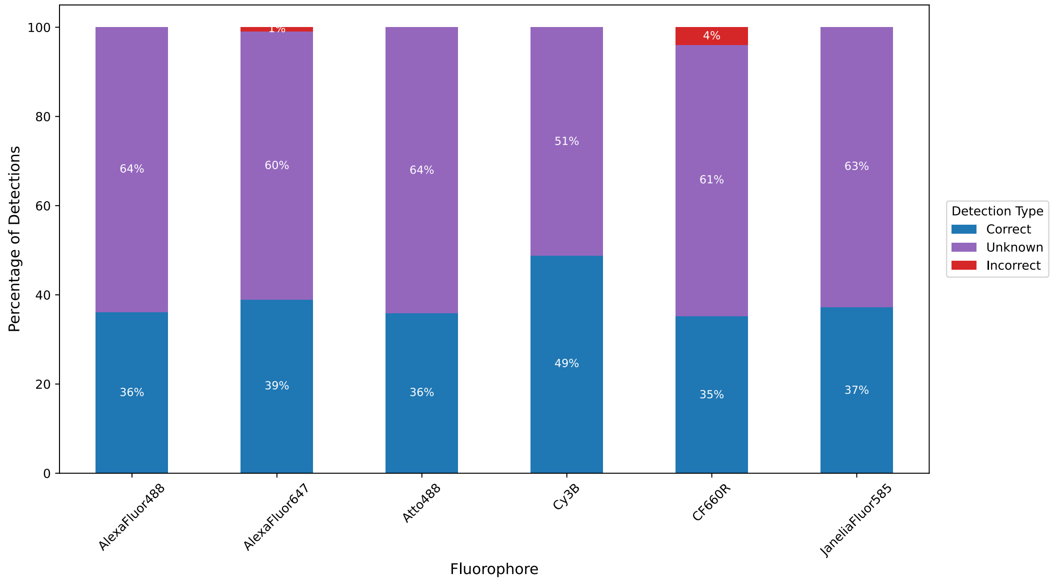


**Figure S1** Percentage of correct, incorrect, and unknown detections using an RGB camera for 6 dyes with photon numbers reported in[1] and identification region widths of 60%.

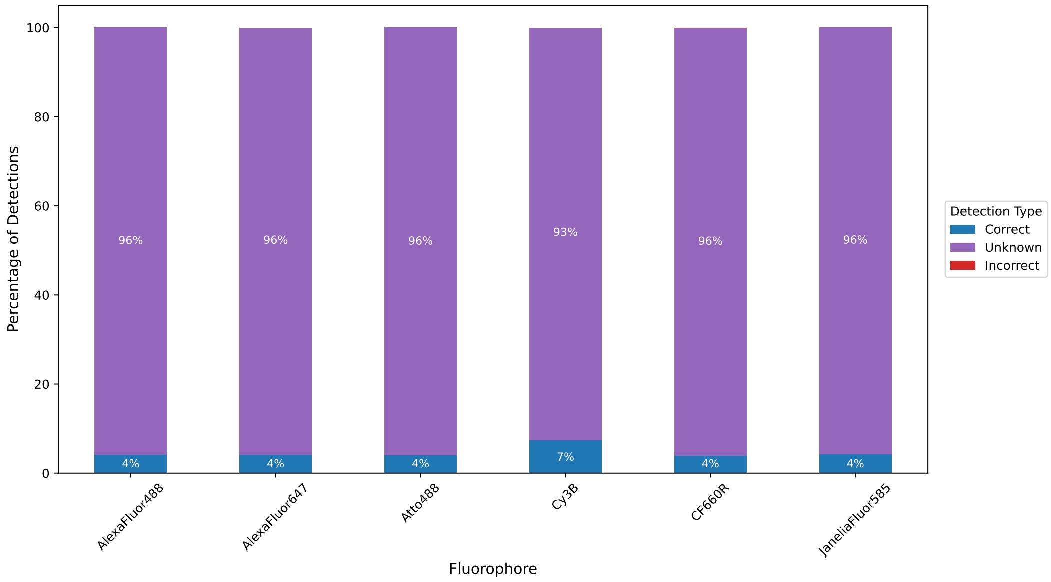


**Figure S2** Percentage of correct, incorrect, and unknown detections using an RGB camera for 6 dyes with photon numbers reported in[1] and identification region widths of 20%.

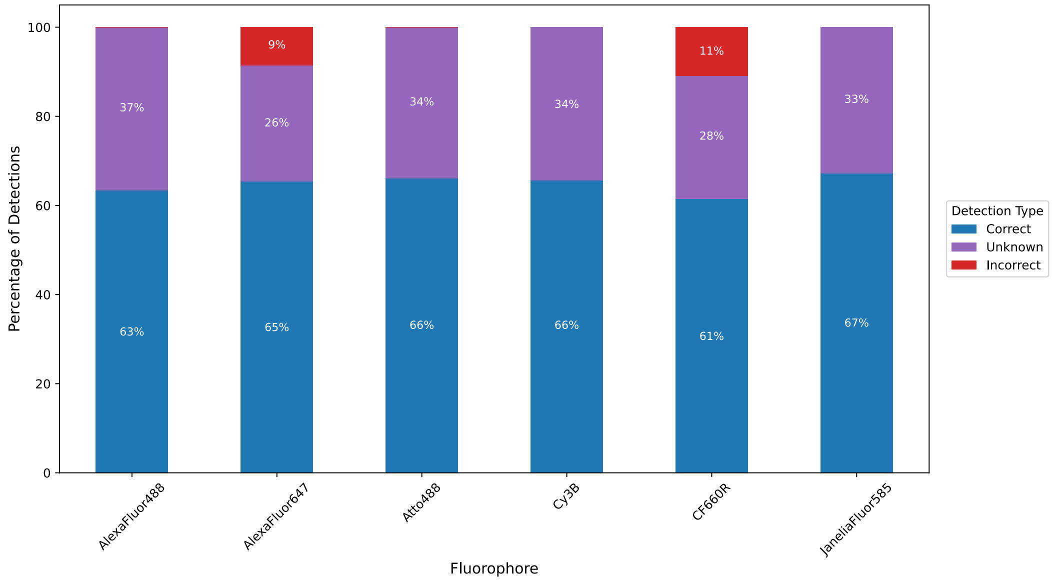


**Figure S3** Percentage of correct, incorrect, and unknown detections using an RGB camera for 6 dyes with photon numbers reported in[1] and identification region widths of 20%.

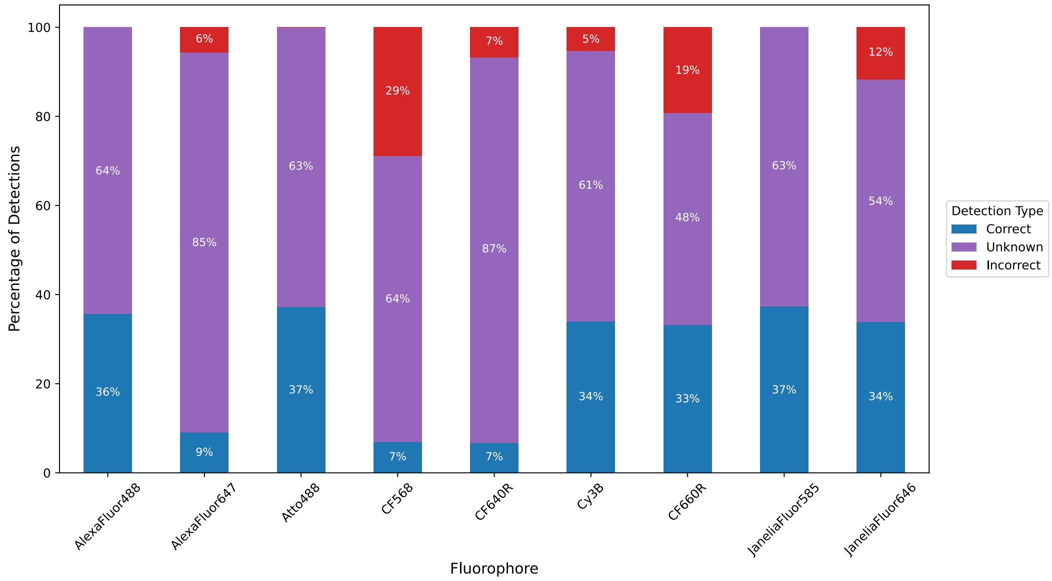


**Figure S4** Percentage of correct, incorrect, and unknown detections using an RGB camera for 9 dyes with photon numbers reported in[1] and identification region widths of 60%.

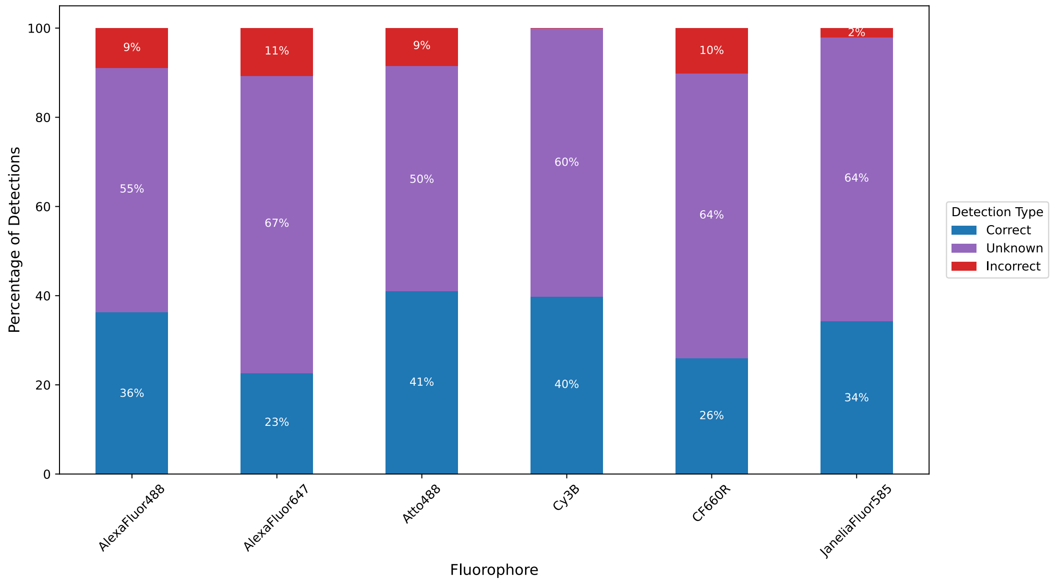


**Figure S5** Percentage of correct, incorrect, and unknown detections using an RGB camera for 6 dyes with 3x reduced photon numbers from originally reported in[1] and identification region widths of 60%.

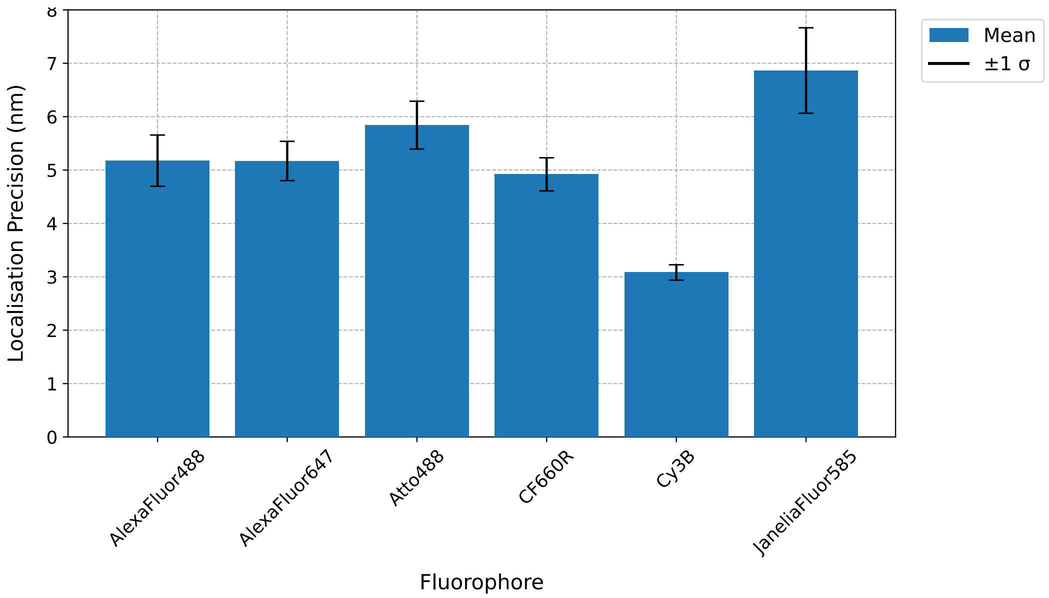


**Figure S6** Localisation precisions with an RGB camera for 6 dyes with 3x reduced photon numbers from originally reported in[1] and identification region widths of 60%.

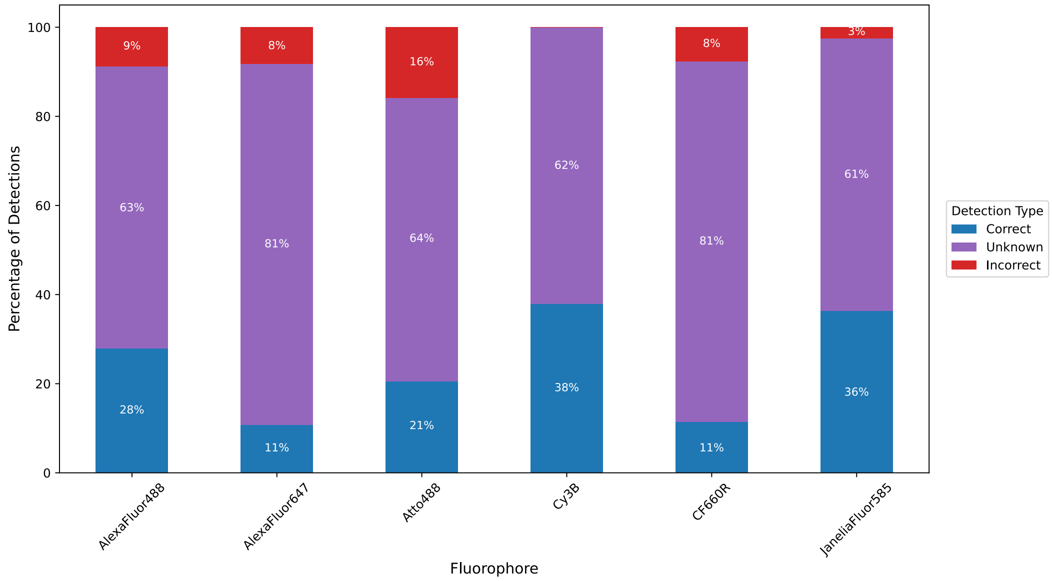


**Figure S7** Percentage of correct, incorrect, and unknown detections using an RGB camera for 6 dyes with 5x reduced photon numbers from originally reported in[1] and identification region widths of 60%.

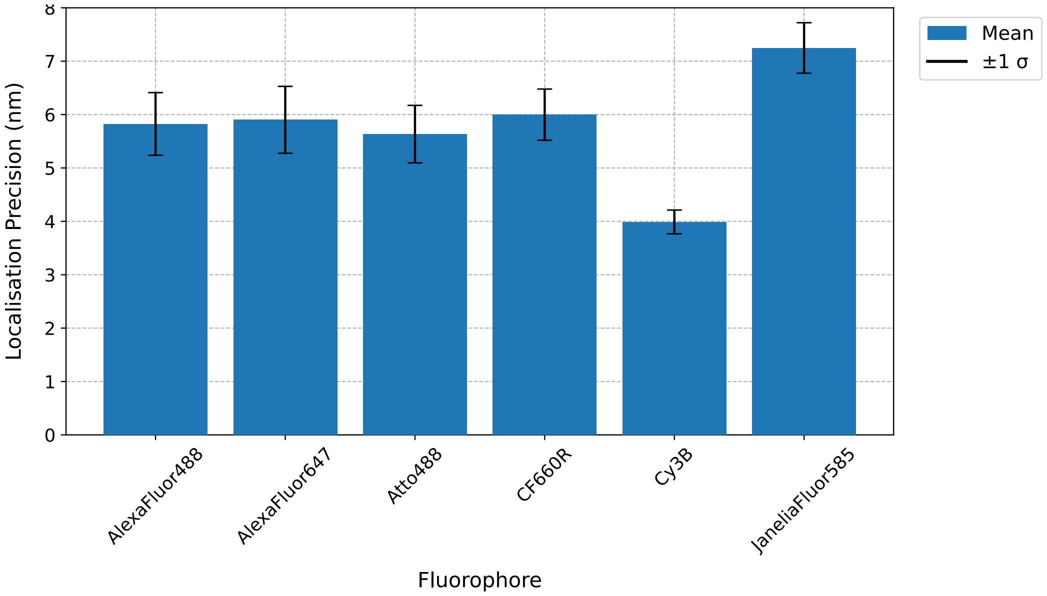


**Figure S8** Localisation precisions with an RGB camera for 6 dyes with 5x reduced photon numbers from originally reported in[1] and identification region widths of 60%.

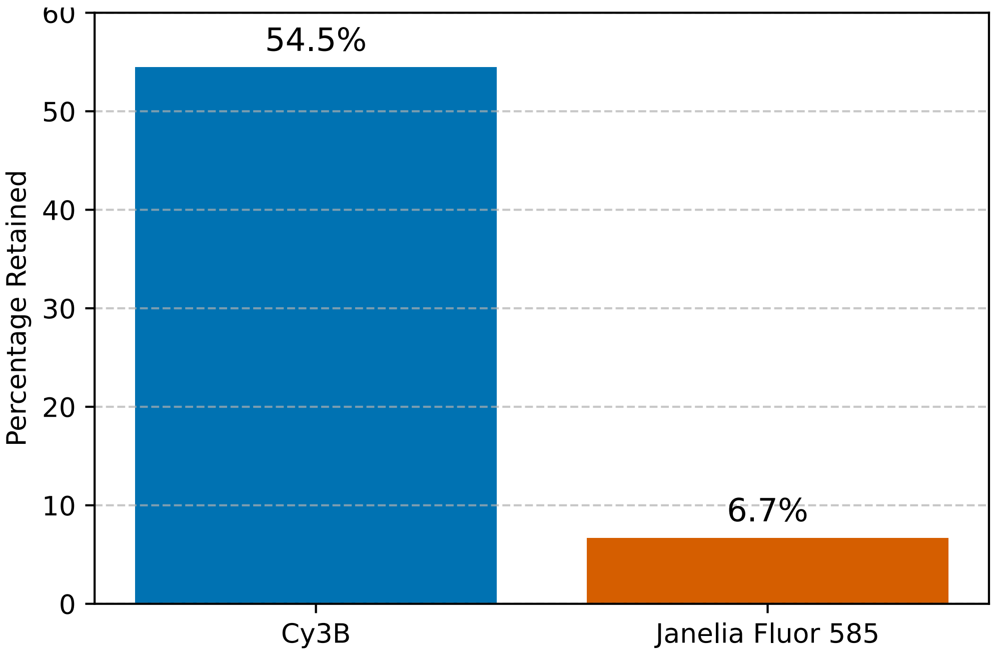


**Figure S9** Percentage of localisations retained after filtering at 5x reduced photon numbers from originally reported in[1]. Total simulated emitters for each dye = 10,000.

### Tables

| Fluorophore | Photons number per 100 ms |
|---|---|
| AlexaFluor 488 | 2811 |
| AlexaFluor 647 | 10348 |
| Atto 488 | 2073 |
| CF568 | 13388 |
| CF640R | 12024 |
| CF660R | 10399 |
| Cy3B | 23195 |
| JaneliaFluor 585 | 2429 |
| JaneliaFluor 646 | 14440 |

**Table S1** Photons emitted per 100 ms under Total Internal Reflection Fluorescence illumination (TIRF) and laser power densities of 100 W/cm2 (488 nm), 175 W/cm2 (561 nm), 350 W/cm2 (638 nm) as experimentally demonstrated in[1].

### References

1. The DNA-PAINT palette: a comprehensive performance analysis of fluorescent dyes | Nature Methods. https://www.nature.com/articles/s41592-024-02374-8.
